# Pathogenic mutations in Clathrin Heavy Chain associated with intellectual disability impair synaptic architecture and learning in *Drosophila*

**DOI:** 10.1101/2025.08.28.672774

**Authors:** Jyoti Das, Manisha Datta, Srikanth Pippadpally, Bhavika Jethani, Surya Bansi Singh, Akankhya Jena, Vimlesh Kumar, Amitabha Majumdar, Deepa Subramanyam

**Author notes:** to whom correspondence should be addressed: Deepa Subramanyam. these authors contributed equally (Manisha Datta, Srikanth Pippadpally, Bhavika Jethani).

## Abstract

Clathrin-mediated endocytosis is essential for neural development and function. Recent studies have linked *de novo* mutations in the clathrin heavy chain (Chc) gene to a range of neurodevelopmental disorders. In this study we have modelled two pathogenic mutations: L1047P and W1108R in *Drosophila melanogaster* and examined their effects on vesicle dynamics, ligand uptake, neuronal development and memory formation. Our data shows that expression of these mutant forms of Chc result in reduced survival and defective learning when expressed ubiquitously or exclusively in neurons. Our analysis also reveals that these mutations have the ability to disrupt vesicle dynamics and reduce ligand uptake in cells. Although we do not see a defect in neuronal morphology and function at the larval neuro-muscular junction, we see an increase in the number of Dlg-negative boutons, and a significant reduction in Spectrin expression, indicative of disruptions in the process of synapse maturation. Overall, this study provides mechanistic insights into the cellular and molecular basis of Chc-related neurodevelopmental disorders.

## Introduction

Neurodevelopmental disorders (NDDs) are a group of disorders characterized by developmental deficits in cognition, and/or motor skills. These include intellectual disability (ID), autism spectrum disorder, attention deficit hyperactivity disorder (ADHD), and speech and language impairment (Khodosevich & Sellgren, 2023). Although factors such as dysregulation in maternal hormones, exposure of the mother to toxins and infections and foetal and neonatal trauma can result in intellectual disability, most NDDs have been shown to have genetic underpinnings. One gene recently associated with ID and global developmental delay (GDD) is the CHC17 gene, coding for the clathrin heavy chain protein. *De novo* mutations in this gene are associated with variable phenotypes such as ID ranging from mild to severe, craniofacial dysmorphisms such as long palpebral fissures and midface hypoplasia, neurological phenotypes such as ataxia and hypotonia, structural brain abnormalities such as hypoplasia of the corpus callosum, microcephaly and epileptic seizures(Demari et al., 2016; Hamdan et al., 2017; Manti et al., 2019; Nabais Sá et al., 2020). Other phenotypes include skeletal abnormalities such as spina bifida occulta and mid-rib hypoplasia (Martín Fernández-Mayoralas et al., 2021), respiratory abnormalities and abnormal kidney function (Itai et al., 2022).

Prior studies have shown that patients harbouring mis-sense and in-frame variants of CHC17 exhibit more severe phenotypes in comparison to individuals carrying frameshift variants. These observations led investigators to hypothesize that mis-sense and in-frame mutations show dominant-negative effects whereas the frameshift mutations result in nonsense-mediated mRNA decay which ultimately leads to haploinsufficiency (Nabais Sá et al., 2020). However, patients with in-frame mutant variants show variable phenotypes. For example, the recurrent c.2669C > T (p.P890L; NM_004859.4) mutation is associated with mild to moderate ID, whereas the c.3140 T > C (p.L1047P) mutation is associated with severe ID, structural abnormalities in the brain and epileptic seizures (Nabais Sá et al., 2020), indicating that different mutations may affect neural development and function in distinct ways. Given that these mutations have only recently been reported, the underlying mechanisms that lead to such a broad spectrum of phenotypes are largely unknown.

CHC17 encodes the clathrin heavy chain, a key component of the clathrin coat best known to carry out the process of receptor-mediated endocytosis in cells. Three heavy chains, each associated with a clathrin light chain forms a clathrin triskelion which polymerizes onto the inner leaflet of the budding plasma membrane to form clathrin-coated pits (CCPs) and drive the endocytosis of ligand-bound receptors present on the plasma membrane (Briant et al., 2020; Kaksonen & Roux, 2018). Since this process spatially and temporally regulates the receptors on the plasma membrane, it can regulate signalling which, when gone awry, can lead to severe developmental defects (Bökel & Brand, 2014; Narayana et al., 2019; Sharma et al., 2025; Tiwari et al., 2025). Apart from endocytosis, clathrin is also known to regulate synaptic vesicle recycling, a process critical for efficient transmission across the synapse (Heerssen et al., 2008; Kasprowicz et al., 2008; Redlingshöfer et al., 2020). Clathrin has also been implicated in the uptake of neurotransmitter receptors, regulating their distribution at the synapse (Ehlers, 2000; Hanley, 2018; Sathler et al., 2021). Given these established roles, it can be expected that mutations in the genes coding for the clathrin heavy chain would show severe neurodevelopmental phenotypes. A previous study in *C.elegans* has also shown that the P892L substitution (P890L mutation in humans) results in defects in synaptic transmission indicating that the process of synaptic vesicle recycling is hampered in the presence of mutant forms of clathrin (Pannone et al., 2023).

The use of *Drosophila melanogaster* as a system to study ID and other NDDs is well appreciated (Androschuk et al., 2015; Coll-Tane et al., 2019). It is a powerful system used to study brain development, synapse function, plasticity and behaviour. Further, the high degree of conservation of the sequence of the clathrin heavy chain protein, and the pathway thereof from humans to *Drosophila*, permits utilization of a variety of model systems to dissect the effect of pathogenic mutations in the context of development. In this study, we have selected two of the pathological missense mutations (c.3140T>C, p.L1047P and c.3322T>C, p.W1108R) and modelled them in *Drosophila melanogaster* to understand how these mutations affect intracellular trafficking, neural transmission and ultimately behaviour. Here, we generated and characterized *Drosophila* lines overexpressing either the wild type or the pathological variants of clathrin tagged to the fluorescent protein, GFP. We show that, the mutant forms of clathrin alters vesicle dynamics, ligand uptake and synaptic architecture. Moreover, *Drosophila* lines carrying these mutations exhibit structural abnormalities in the brain and display defects in learning and memory. These phenotypes mimic the abnormalities observed in the human patients, thereby validating the utility of this model for studying pathological clathrin mutations, while providing a system to explore the molecular underpinnings of this disorder.

## Results

### Overexpression of pathogenic CHC variants in *Drosophila* reduces survival and alters intracellular trafficking

We began our study by selecting point mutations in the CHC 17 gene that resulted in intellectual disability. Since the missense and in-frame variants showed a more severe phenotype in humans, we decided to model these mutations in *Drosophila*. Given the highly conserved nature of the CHC protein (human CHC shares 99% sequence similarity with mouse *Chc* and 80% sequence similarity with *Drosophila* Chc), we observed that most point mutations reported in humans could be mapped onto the *Drosophila* protein sequence with a shift of one amino acid (Supplementary Fig. 1A). We selected two point mutations – L1047P and W1108R (L1048P and W1109R in *Drosophila*) and generated fly lines overexpressing either GFP-tagged wild type Chc or either of the two mutants under a UAS-promoter (Supplementary Fig.1B). Since the L1047P-Chc mutation is an extremely severe mutation, and has been shown to result in lethality in the homozygous condition in *C. elegans* (Pannone et al., 2023), we suspected that an attempt to generate L1047P-Chc mutant flies without the endogenous Chc may also result in lethality at early developmental stages. Therefore, we did not alter the endogenous Chc locus, but simply overexpressed the mutant forms of Chc using the GAL4-UAS system.

To begin the characterization of these flies, we first crossed them to a tubulin GAL4 to drive their expression ubiquitously in all tissues. We observed that the animals expressing the L1048P-Chc mutant were lethal at early larval stages. Contrary to L1048P-Chc mutants, the W1109R-Chc mutants were able to reach adulthood and the eclosion percentage of W1109R-Chc mutants was comparable to wild type Chc overexpressing flies (Fig.1A). When the L1048P-Chc was expressed using a pan-neuronal driver (Elav-GAL4), the number of eclosed adults was significantly lower than in wild-type expressing flies (Fig.1B, Supplementary Fig.1C) indicating that even in *Drosophila*, the L1047P mutation is more severe compared to the W1108R mutation. We then studied the survival of these mutant flies and observed that ubiquitous (Fig.1C, Supplementary Fig. 1D) or pan-neuronal overexpression (Fig.1D, Supplementary Fig.1E) of these point mutations led to a significant reduction in their survival compared to the controls, indicating hitherto unknown deleterious effects of these pathogenic Chc mutations.

**Figure 1:**
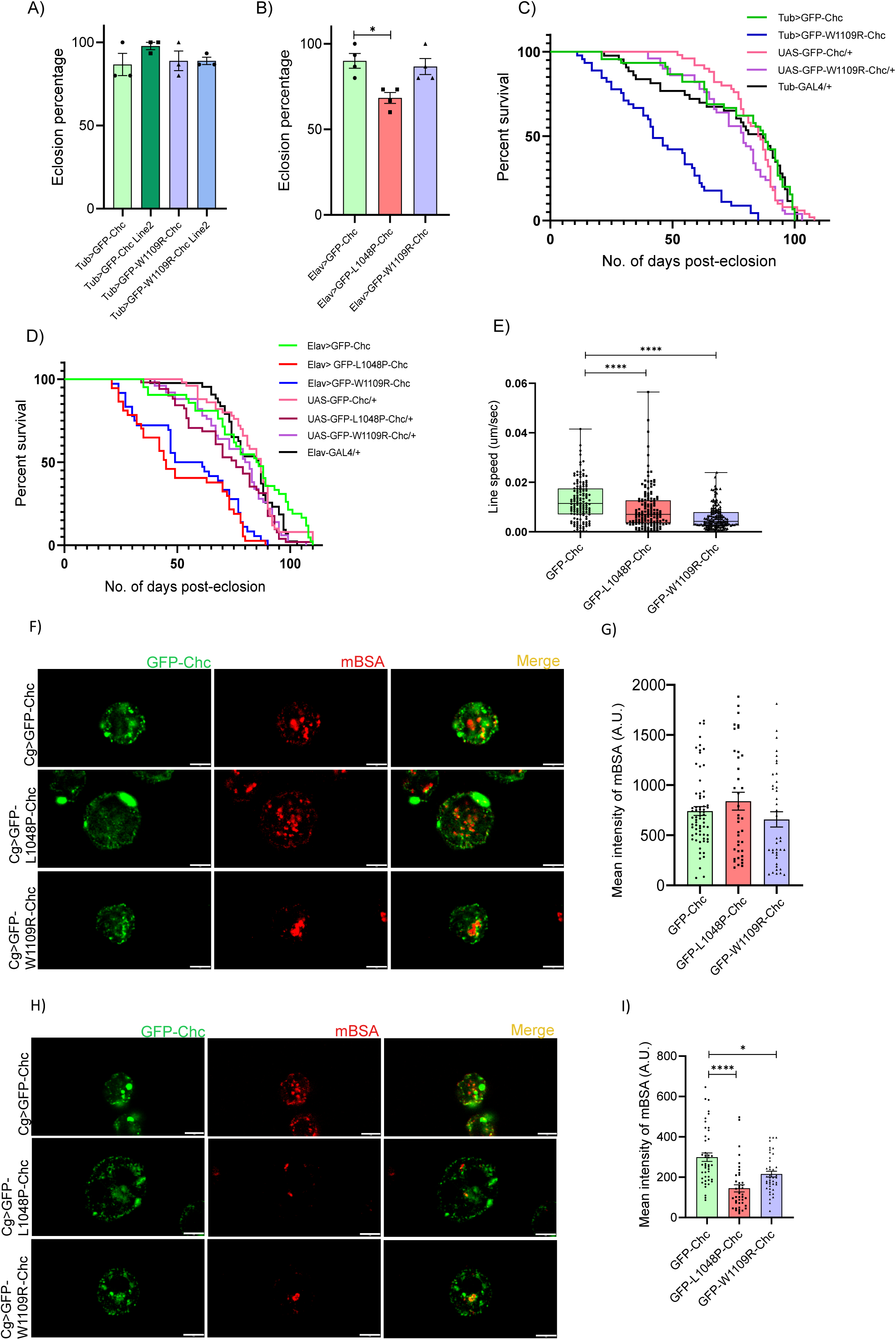
Pathogenic mutants of Chc affect survival and clathrin-mediated endocytosis. **A)** Eclosion percentage of *Tub>GFP-W1109R-Chc* (Line1 and Line2) compared to *Tub>GFP-Chc* (Line1 and Line2). n=45; N=3. Statistical significance calculated using Kruskal-Wallis test for non-parametric data. **B)** Eclosion percentage of *Elav>GFP-L1048P-Chc* and *Elav>GFP-W1109R-Chc* compared to *Elav>GFP-Chc*. n=60; N=4. P value calculated using the Kruskal-Wallis test for non-parametric data. *p<0.05. **C)** Kaplan-Meier curve depicting survival of *Tub>GFP-W1109R-Chc* (N=45) flies compared to *Tub>GFP-Chc* (N=45), *UAS-GFP-W1109R-Chc/+*(N=50), *UAS-GFP-Chc/+* (N=50) and *Tub-GAL4/+*(N=43) controls. **D)** Kaplan-Meier curve depicting survival of *Elav>GFP-W1109R-Chc* (N=36) and *Elav>GFP-L1048P-Chc* (N=37) flies compared to *Elav>GFP-Chc* (N=42), *UAS-GFP-W1109R-Chc/+* (N=50), *UAS-GFP-L1048P-Chc/+* (N=51) *UAS-GFP-Chc/+* (N=50) and *Elav-GAL4/+* (N=43) controls. **E)** Graph representing line speed of vesicles in hemocytes expressing *GFP-Chc* (n=137), *GFP-L1048P-Chc* (n=150) and *GFP-W1109R-Chc* (n=152). P-value calculated using Kruskal-Wallis test for non-parametric data. ****P≤0.0001. **F)** Representative images of mBSA uptake in *GFP-Chc*, *GFP-L1048P-Chc* and *GFP-W1109R-Chc* expressing hemocytes. Ligand uptake was performed at 23°C. **G)** Graph quantifying the mBSA intensity inside hemocytes expressing *GFP-Chc* (n=69), *GFP-L1048P-Chc* (n=41) and *GFP-W1109R-Chc* (n=40). Ligand uptake was performed at 23°C. Statistical significance was calculated using the Kruskal-Wallis test for non-parametric data. **H)** Representative images of mBSA uptake in *GFP-Chc*, *GFP-L1048P-Chc* and *GFP-W1109R-Chc* expressing hemocytes. Ligand uptake was performed at 37°C. **I)** Graph quantifying the mBSA intensity inside hemocytes expressing *GFP-Chc* (n=45), *GFP-L1048P-Chc* (n=41) and *GFP-W1109R-Chc* (n=48). Ligand uptake was performed at 37°C. Statistical significance calculated using the Kruskal-Wallis test for non-parametric data. *P≤0.05; ****P≤0.0001. All data is represented as Mean ± SEM. Scale bar – 5µm.

Given that the primary role of the clathrin heavy chain is to regulate uptake of ligand-bound receptors from the plasma membrane by forming clathrin-coated vesicles (CCVs), we asked whether this function was affected in the presence of these mutants. To study this, we isolated hemocytes from *Drosophila* larvae expressing these mutants under the Cg-GAL4 promoter, and studied the dynamics of CCVs using live cell imaging. We observed that the movement of vesicles overexpressing either L1048P-Chc or W1109R-Chc was significantly reduced compared to vesicles overexpressing WT-Chc as seen by the difference in the line speed of the vesicles (Fig.1E). To confirm that this phenotype was also observed in a mammalian system, we generated these point mutations in the human Chc ORF, using site directed mutagenesis (Supplementary Fig. 1F, G) and overexpressed them in HEK-293 cells to study vesicle dynamics. We saw that overexpression of L1047P-Chc did not result in the formation of distinct vesicles but exhibited a diffused expression pattern (Supplementary Fig.1H) due to which vesicles could not be identified and the line speed could not be calculated. We therefore estimated the line speed for W1108R-Chc vesicles and observed a significant reduction in their line speed compared to WT-Chc vesicles (Supplementary Fig.1I).

As the expression of mutant forms of Chc resulted in a reduced speed of CCVs, we speculated that ligand uptake may also be compromised. We quantitated ligand uptake in the presence of mutant forms of Chc by studying the uptake of maleylated BSA in *Drosophila* hemocytes. We did not observe any defect in the total ligand uptake in the cells expressing mutant forms of Chc, possibly due to the presence of endogenous wild type Chc (Fig. 1F, G). We therefore reasoned that if there were subtle differences in ligand uptake in the presence of the mutant, these would be exaggerated in the presence of a cellular stress, which would affect the rate of endocytosis. Given that the process of clathrin-mediated endocytosis is temperature sensitive (Tomoda et al., 1989; Weigel & Oka, 1981), we elevated the temperature to assay its effect on ligand uptake. At 37°C, we observed that the total ligand uptake in cells overexpressing L1048P-CHC and W1109R-CHC was lower compared to the cells overexpressing wild type CHC (Fig. 1H, I, Supplementary Fig. 1J, K).

Overall, our data suggests that the L1048P-Chc and W1109R-Chc point mutations alter intracellular trafficking, reduce eclosion of adults from pupae and significantly reduce survival. Our data also suggests that compared to the W1109R-CHC mutant, the L1048P-CHC mutant is much more debilitating, given that its overexpression results in lethality at early developmental stages.

### Mutant forms of Chc do not show functional defects at the larval neuromuscular junction

Given that the role of clathrin in synaptic vesicle recycling at the *Drosophila* larval neuromuscular junction (NMJ) is well established (Heerssen et al., 2008; Kasprowicz et al., 2008), we asked whether the presence of mutant forms of Chc resulted in functional defects at the NMJ. Electrophysiology studies did not show any differences in mEJP frequency (Fig. 2A, Supplementary Fig. 2A), amplitude (Fig.2B, Supplementary Fig. 2B), EJP amplitude (Fig.2C, Supplementary Fig. 2C) or quantal content (Fig.2D, Supplementary Fig. 2D), indicating that the presence of Chc mutants did not affect synaptic vesicle recycling at the larval NMJ. This was further corroborated by the fact that there was no difference in larval locomotion between larvae expressing WT and mutant forms of Chc (Fig.2E).

**Figure 2:**
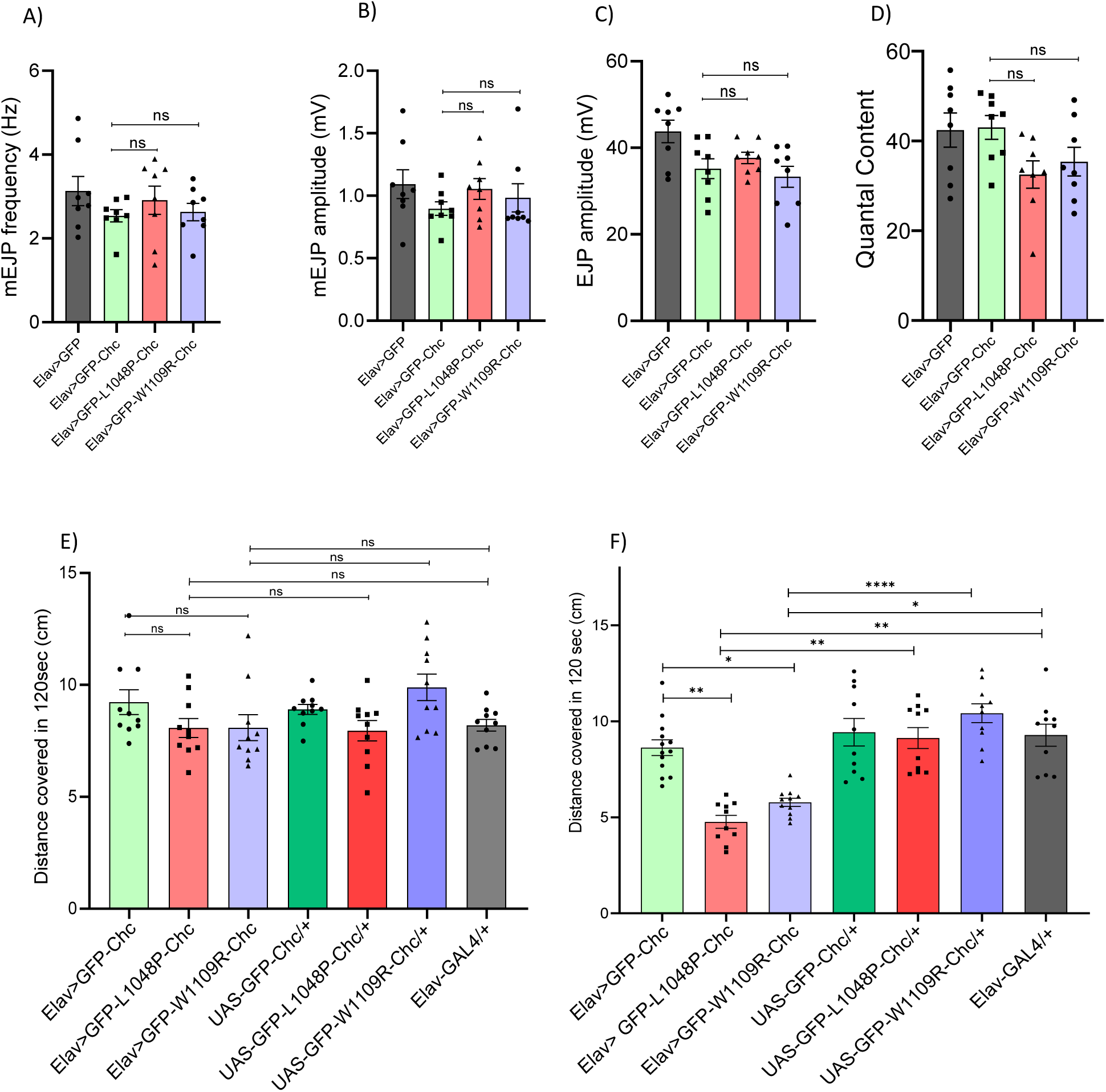
Mutant forms of Chc do not show functional defects at the larval neuromuscular junction **A)** Histograms showing the mEJP frequency in *Elav-Gal4>UAS-GFP* (3.13 ± 0.34), *Elav>GFP-Chc* (2.54 ± 0.14), *Elav>GFP-L1048P-Chc* (2.91 ± 0.33), and Elav>ChcW1109R (6959) (2.63 ± 0.20). Statistical analysis is based on one-way ANOVA followed by post-hoc Tukey’s multiple-comparison test. 8 animals were analyzed for each genotype. **B)** Histograms showing average mEJP amplitude in *Elav-Gal4>UAS-GFP* (1.09 ± 0.11), *Elav>GFP-Chc*(0.89 ± 0.05), *Elav>GFP-L1048P-Chc* (1.05 ± 0.08), and *Elav>GFP-W1109R-Chc* (0.98 ± 0.11). Statistical analysis is based on one-way ANOVA followed by post-hoc Tukey’s multiple-comparison test. 8 animals were analyzed for each genotype. **C)** Histograms showing average EJP amplitude in *Elav-Gal4>UAS-GFP* (43.8 ± 2.60), *Elav>GFP-Chc* (35.2 ± 2.30), *Elav>GFP-L1048P-Chc* (37.7 ± 1.32), and *Elav>GFP-W1109R-Chc* (33.3 ± 2.44). Statistical analysis is based on one-way ANOVA followed by post-hoc Tukey’s multiple-comparison test. 8 animals were analyzed for each genotype. **D)** Histograms showing quantal content in *Elav>GFP* (42.4 ± 3.79), *Elav>GFP-Chc* (43.0 ± 2.65), *Elav>GFP-L1048P-Chc* (32.5 ± 3.04), and *Elav>GFP-W1109R-Chc* (35.4 ± 3.17). Statistical analysis is based on one-way ANOVA followed by post-hoc Tukey’s multiple-comparison test. 8 animals were analyzed for each genotype. **E)** Graph representing the distance travelled by *Elav>GFP-W1109R-Chc*, *Elav>GFP-L1048P-Chc* larvae compared to controls *Elav>GFP-Chc*, *UAS-GFP-W1109R-Chc/+*, *UAS-GFP-L1048P-Chc/+*, *UAS-GFP-Chc/+* and *Elav-GAL4/+*. Statistical significance was calculated using one-way ANOVA. 10 animals of each genotype were studied. The larvae were kept at 25°C. **F)** Graph representing the distance travelled by Elav>GFP-W1109R-Chc, Elav>GFP-L1048P-Chc larvae compared to controls *Elav>GFP-Chc*, *UAS-GFP-W1109R-Chc/+*, *UAS-GFP-L1048P-Chc/+*, *UAS-GFP-Chc/+* and *Elav-GAL4/+*. Statistical significance calculated using one-way ANOVA. *P≤0.05; **P≤0.005; ***P≤0.0005, ****P≤0.0001. 10 animals of each genotype were studied. The larvae were kept at 37°C for 20 minutes. All data is represented as mean ± SEM.

We suspected that the absence of the phenotype may be due to the presence of the endogenous wild type Chc, and that subjecting these larvae to heat stress may reveal a phenotype, in a manner similar to what was observed in the context of ligand uptake. To test this, we studied larval locomotion after subjecting the larvae to heat stress. We observed that the distance travelled by the larvae expressing L1048P-Chc and W1109R-Chc was significantly less compared to WT Chc larvae (Fig.2F, Supplementary Fig. 2E), indicating that the presence of the endogenous Chc may permit normal functioning of endocytosis under non-stressed conditions.

### Neuronal morphology is unaltered in the presence of pathogenic Chc variants

The loss of critical endocytic genes has been associated with altered neuronal morphology at the NMJ (Koh et al., 2004; O’Connor-Giles et al., 2008). Studies have shown that mutants affecting key endocytic proteins lead to changes in synaptic bouton structure and active zone numbers (Dickman et al., 2006; O’Connor-Giles et al., 2008). We asked if similar defects were observed in the presence of pathogenic Chc mutants. To test this, we studied neuronal morphology at the third instar larval NMJ (Fig.3A, Supplementary Fig. 3A). Our study did not reveal any gross differences in neuronal morphology, with total bouton number/NMJ (Fig.3B, Supplementary Fig. 3B), average bouton size (Fig.3C, Supplementary Fig. 3C) and the number of branches/NMJ (Fig. 3D, Supplementary Fig. 3D) being comparable to that of wild type expressing lines.

**Figure 3:**
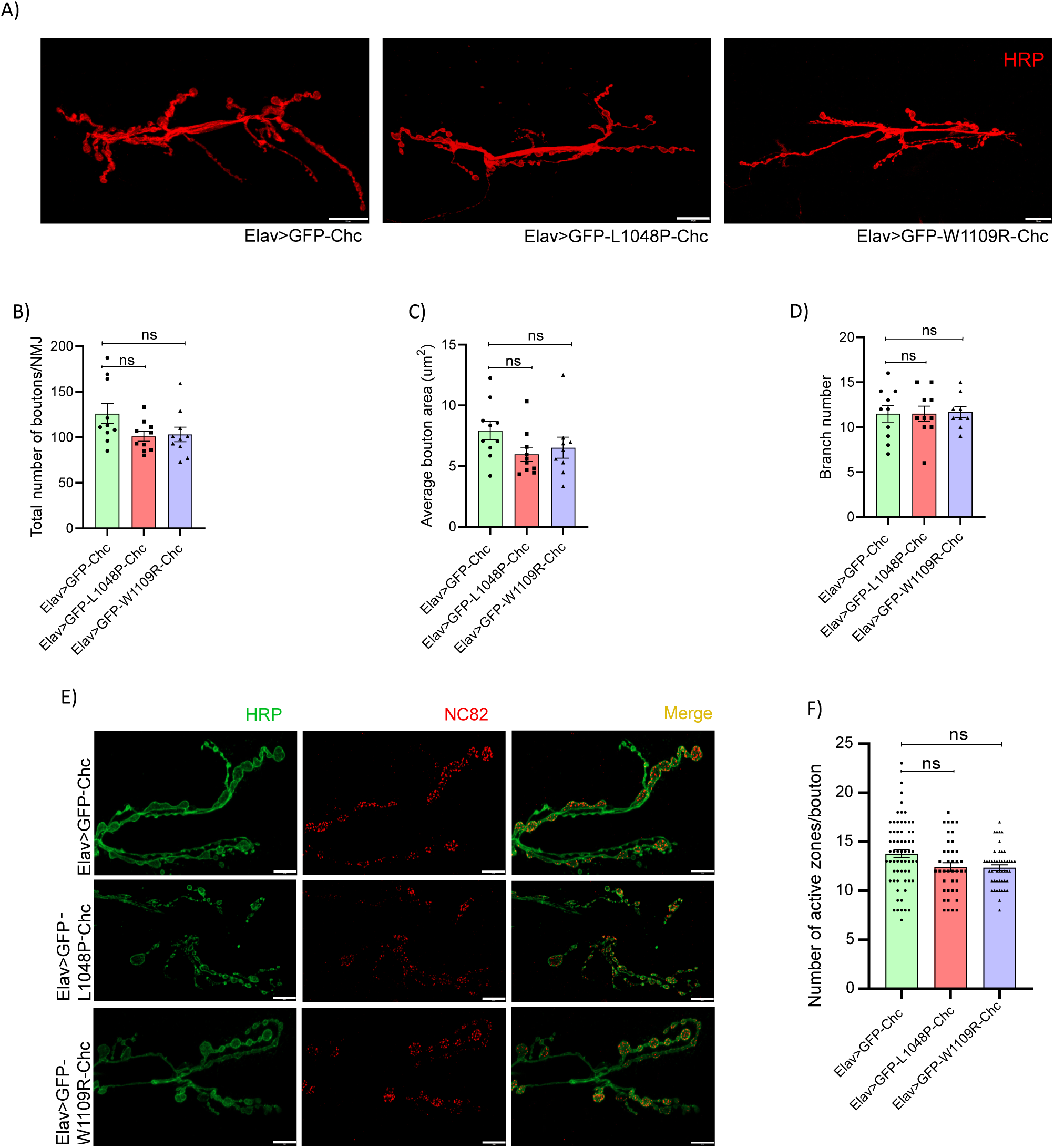
Neuronal morphology is unaltered in the presence of pathogenic Chc variants **A)** Representative images of neuronal morphology in *Elav>GFP-Chc*, *Elav>GFP-L1048P-Chc* and *Elav>GFP-W1109R-Chc* larvae. Neurons at the A2/A3 hemisegment on the 6th/7th muscle were imaged. 5-7 animals were imaged for each genotype. Scale bar – 20µm. **B)** Graph representing total bouton number in *Elav>GFP-Chc*, *Elav>GFP-L1048P-Chc* and *Elav>GFP-W1109R-Chc* larvae. Statistical significance was calculated using Kruskal-Wallis test. **C)** Graph representing average bouton size in *Elav>GFP-Chc*, *Elav>GFP-L1048P-Chc* and *Elav>GFP-W1109R-Chc* larvae. Statistical significance calculated using the Kruskal-Wallis test. **D)** Graph representing total branch number in *Elav>GFP-Chc* Line2, *Elav>GFP-L1048P-Chc* Line2 and *Elav>GFP-W1109R-Chc* Line2 larvae. Statistical significance calculated using the Kruskal-Wallis test. **E)** Representative images showing the active zones in *Elav>GFP-Chc*, *Elav>GFP-L1048P-Chc* and *Elav>GFP-W1109R-Chc* larvae. Scale Bar – 10µm. **F)** Graph representing the number of active zones per bouton in *Elav>GFP-Chc*, *Elav>GFP-L1048P-Chc* and *Elav>GFP-W1109R-Chc* larvae. 40-90 boutons were counted for each genotype. Statistical significance calculated using the Kruskal-Wallis test. All data is represented as mean ± SEM.

Furthermore, analysis of the active zones in these mutants showed that the total number of active zones/ bouton in larvae expressing L1048P-Chc and W1109R-Chc was comparable to larvae expressing WT-Chc (Fig. 3E, F; Supplementary Fig. 3E, F).

Overall, our results indicate that early development of neuronal structures occurs normally even in the presence of mutant forms of Chc.

### Presence of pathogenic Chc variants alters the postsynaptic architecture

The process of synapse development and maturation is highly coordinated, involving the assembly of active zone proteins at the presynaptic terminal, followed by assembly of the postsynaptic complexes (Chou et al., 2020). The assembly of postsynaptic scaffolding and cytoskeletal proteins and the process of synaptic maturation have been shown to be dependent on the endocytosis of the Frizzled receptor, DFz2 upon binding of Wingless (Wg) at the *Drosophila* NMJ (Ataman et al., 2008; Mosca & Schwarz, 2010), in a dynamin-dependent manner (Mathew et al., 2005). We therefore sought to determine whether the presence of mutant forms of clathrin disrupted the process of synaptic maturation in *Drosophila*.

To assess impaired synaptic maturation, we used two parameters described in a previous study (Restrepo et al., 2022): the presence of HRPL, DlgL boutons (ghost boutons), and a reduction in the postsynaptic cytoskeletal protein, Spectrin. Using these criteria, we examined synapse maturation in larvae overexpressing mutant forms of Chc.

We observed that overexpression of both, L1048P-Chc and W1109R-Chc in the muscle (postsynaptic cell) resulted in an increase in the number of Dlg-negative boutons (Fig.4 A,B; Supplementary Fig. 4A, B) and led to a reduction in the levels of Spectrin at the synapse (Fig.4 C,D; Supplementary Fig. C, D), indicating that the presence of mutant forms of clathrin lead to impaired synaptic maturation. Surprisingly, overexpression of the mutants in the neurons (presynaptic cell) gave us different results for the two mutants. Overexpression of W1109R-Chc in the neurons did not affect synapse maturation as expected, but overexpression of L1048P-Chc led to an increase in the number of Dlg-negative boutons (Fig.4E, F; Supplementary Fig. 4E, F) and a decrease in the levels of Spectrin (Fig.4G, H; Supplementary Fig. 4G, H) at the synapse. It has been suggested that the formation of postsynaptic cytomatrix structures depends on signals from the presynaptic cell. It is possible that overexpression of L1048P-Chc in the neurons could alter signalling from the presynaptic cell to the postsynaptic cell, resulting in impaired neuronal maturation.

**Figure 4:**
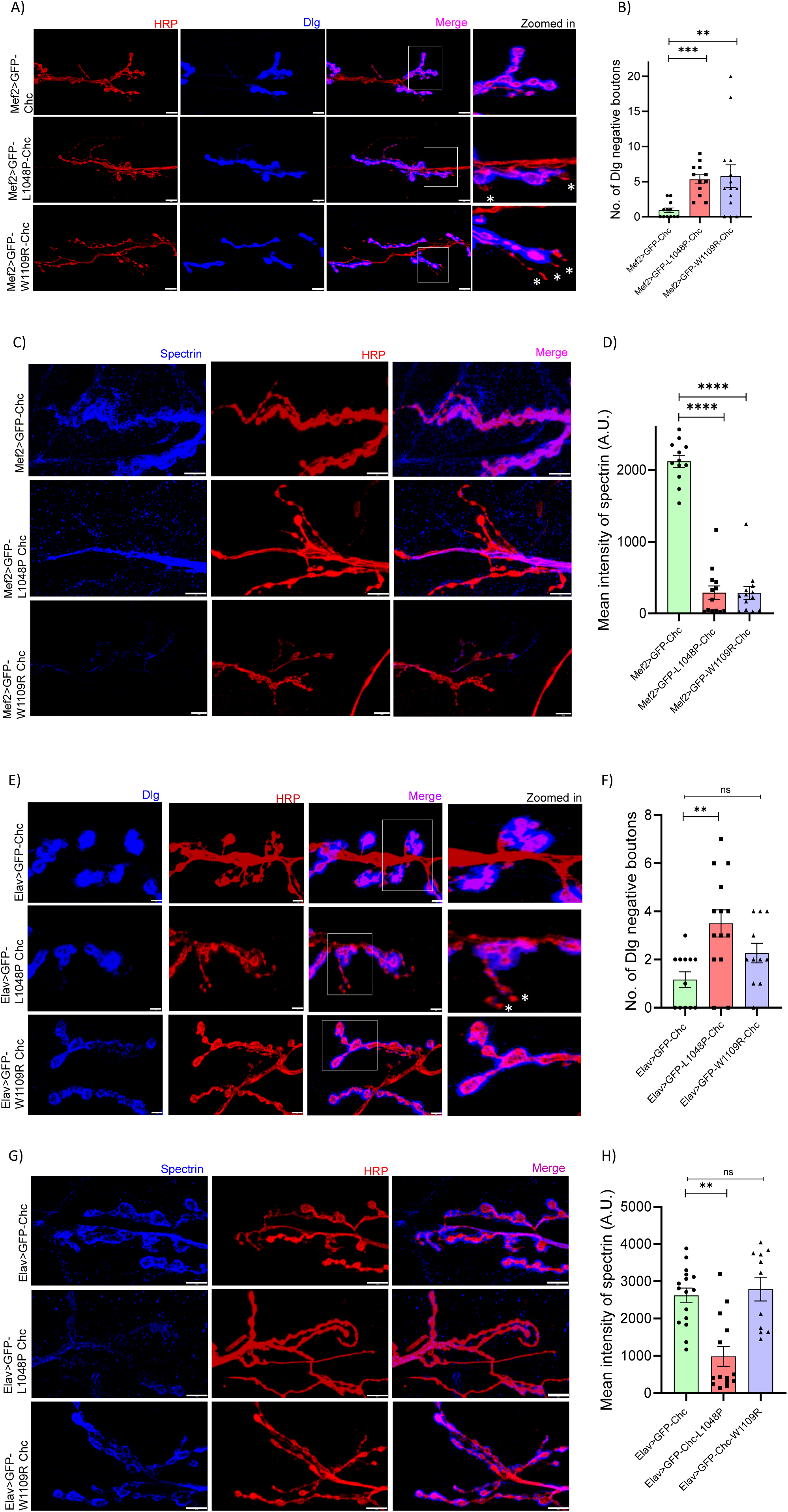
Presence of pathogenic Chc variants alters the postsynaptic architecture **A)** Representative images showing the Dlg positive and negative boutons in *Mef2>GFP-Chc*, *Mef2>GFP-L1048P-Chc* and *Mef2>GFP-W1109R-Chc* larvae. Scale bar-10µm. **B)** Graph quantifying the number of Dlg-negative boutons in *Mef2>GFP-Chc*, *Mef2>GFP-L1048P-Chc* and *Mef2>GFP-W1109R-Chc* larvae. 8-10 animals were dissected for each genotype. Statistical significance calculated using the Kruskal-Wallis test. **P≤0.005; *** P≤0.0005. **C)** Representative images showing Spectrin in *Mef2>GFP-Chc*, *Mef2>GFP-L1048P-Chc* and *Mef2>GFP-W1109R-Chc* larvae. Scale bar-10µm. **D)** Graph quantifying spectrin intensity in *Mef2>GFP-Chc*, *Mef2>GFP-L1048P-Chc* and *Mef2>GFP-W1109R-Chc* larvae. 8-10 animals were dissected for each genotype. Statistical significance calculated using the Kruskal-Wallis test. **** P≤0.0001. **E)** Representative images showing the Dlg-positive and negative boutons in *Elav>GFP-Chc*, *Elav>GFP-L1048P-Chc* and *Elav>GFP-W1109R-Chc* larvae. Scale bar-5µm. **F)** Graph quantifying the number of Dlg negative boutons in Elav>GFP-Chc, Elav>GFP-L1048P-Chc and Elav>GFP-W1109R-Chc larvae. 8-10 animals were dissected for each genotype. Statistical significance calculated using the Kruskal-Wallis test. **P≤0.005. **G)** Representative images showing Spectrin in *Elav>GFP-Chc*, *Elav>GFP-L1048P-Chc* and *Elav>GFP-W1109R-Chc* larvae. Scale bar-10µm. **H)** Graph quantifying spectrin intensity in *Elav>GFP-Chc*, *Elav>GFP-L1048P-Chc* and *Elav>GFP-W1109R-Chc* larvae. 8-10 animals were dissected for each genotype. Statistical significance calculated using the Kruskal-Wallis test. ** P≤0.005. All data is represented as mean ± SEM.

### Defective learning and memory are observed in the *Drosophila* lines expressing pathogenic Chc variants

Patients bearing mutant forms of CHC17 display features of intellectual disability (Hamdan et al., 2017; Manti et al., 2019; Nabais Sá et al., 2020). We asked whether overexpression of these mutant forms of Chc also resulted in behavioural phenotypes similar to humans in *Drosophila*. Given that the primary phenotype observed in humans is intellectual disability, we first studied whether the learning behaviour in flies would be affected upon overexpression of these mutants. To study learning and short-term memory formation we used an aversive taste conditioning paradigm where starved flies were taught to associate sucrose to denatonium benzoate (a bitter substance that flies are aversive to), which would result in a decrease in their proboscis extension response (PER) in response to sucrose (Fig.5A). We observed that overexpression of either L1048P-Chc and W1109R-Chc in the neurons did not lead to a significant reduction in PER in the third training and the testing, indicating that these flies had defects in learning and memory formation (Fig.5B; Supplementary Fig. 5B). We observed similar results upon overexpression of W1109R-Chc ubiquitously using Tubulin-GAL4 (Fig.5C, Supplementary Fig. 5A). Given that gustatory learning in *Drosophila* is dependent on the mushroom body (Kirkhart et al, 2015), we asked whether the architecture of the mushroom body was altered in the presence of these mutants. Analysis of these structures using FasII revealed defects in the mushroom body structures in ∼30% of flies (3 out of 9 brains for L1048P-Chc; and 3 out of 10 brains for W1109R-Chc) studied (Fig.5D). In summary, these results suggest that neuronal expression of Chc point mutations results in impaired memory formation possibly due to defective mushroom body structures, highlighting the neurodevelopmental defects caused by these point mutations.

**Figure 5:**
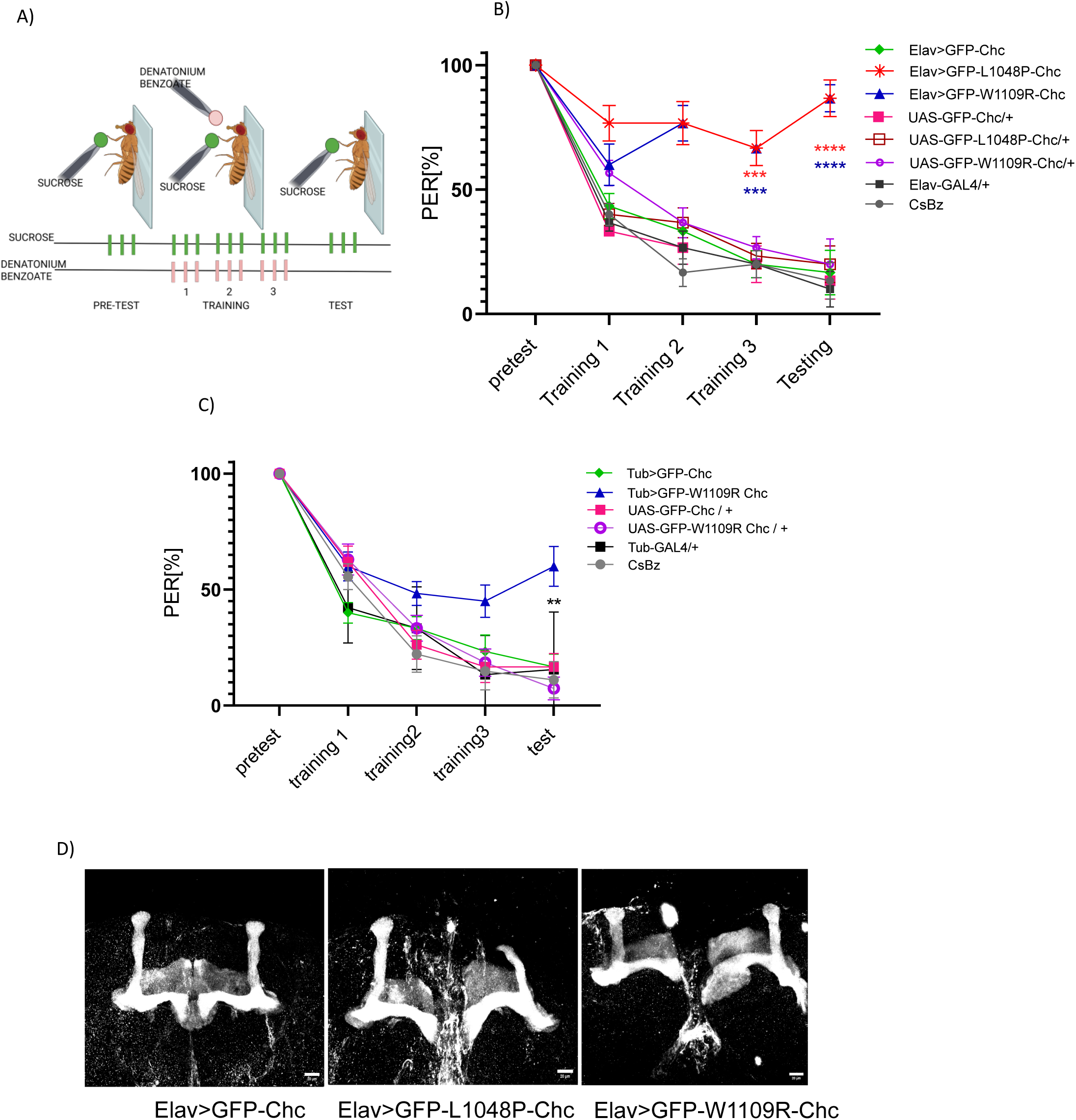
Defective learning and memory are observed in the *Drosophila* lines expressing pathogenic Chc variants **A)** Schematic representation of the proboscis extension response (PER) assay done to assess learning in *Drosophila*. A pretest of stimulation with sucrose alone is followed by three training trial triplets, where sucrose is immediately followed by the application of denatonium benzoate (green) to the extended proboscis. Only sucrose is provided during the test, and PER is measured. **B)** PER assay for *Elav>GFP-L1048P-Chc*, *Elav>GFP-W1109R-Chc* flies and *Elav>GFP-Chc* and parental controls. N=10-20 for each genotype. All flies were 3-10 days old post eclosion. Statistical significance was calculated using a two-way ANOVA followed by the Dunnett multiple comparison method. ***P≤0.0005; ****P≤0.0001. Statistical significance shown using *Elav>GFP-Chc* as a control. **C)** PER assay for *Tub>GFP-W1109R-Chc Line2* flies and *Tub>GFP-Chc* Line2 and parental controls. N=10-20 for each genotype. All flies were 3-10 days old post eclosion. Statistical significance was calculated using a two-way ANOVA followed by the Dunnett multiple comparison method. ** P≤0.005. Statistical significance shown using *Tub>GFP-Chc* Line2 as a control. **D)** Representative images of mushroom bodies in *Elav>GFP-Chc*, *Elav>GFP-L1048P-Chc* and *Elav>GFP-W1109R-Chc* expressing flies.Scale bar-20µm All data is represented as mean ± SEM.

## Discussion

The recently reported mutations in the CHC17 (Chc in *Drosophila*) gene have been associated with a wide range of behavioural phenotypes that include mild to severe ID, microcephaly, epileptic seizures, along with other structural defects in the brain and kidneys. Our work attempts to provide an insight into the mechanistic understanding of how two of the reported mutations - L1047P and W1108R, affect intracellular trafficking and synaptic architecture that ultimately regulates behaviour and survival.

We show that overexpression of mutant forms of Chc affects vesicle dynamics and ligand uptake, indicating that these point mutations may alter properties such as triskelion stability, light chain binding, and rate of polymerization. Further assays aimed at understanding how these mutations alter the biochemical properties of clathrin would provide valuable insights into whether established clathrin-dependent processes, such as ligand-bound receptor endocytosis and synaptic vesicle recycling, are altered in the presence of these mutants. This may also explain why the L1047P-Chc mutant displays more severe defects than the W1108R-Chc mutant in humans and *Drosophila*.

While we expected to find defects in neuronal morphology, we were surprised to find that the presence of these mutants led to disruption of the postsynaptic cytomatrix. As described earlier, our data suggests that the process of neuronal maturation is impaired in the presence of these mutants. Previous studies (Mathew et al, 2005; Mosca and Shwartz,2010; and Restrepo et al, 2022) show that the process of synapse maturation at the *Drosophila* NMJ is dependent on a non-canonical Wnt signaling pathway requiring the endocytosis of the Wingless (Wg) bound Frizzled receptor (DFz2), the C-terminal of which is then cleaved and trafficked into the nucleus to regulate the expression of genes involved in synaptic maturation. Mathew et al, (2005) also showed that the trafficking of the cleaved C-terminal to the nucleus is hampered in *shibire* (*Drosophila* ortholog of Dynamin) mutants, indicating that the endocytosis of Wg-bound DFz2 may likely be clathrin and dynamin dependent. Furthermore, the endocytosis of the DFz2 receptor from the basolateral surface of the developing wing imaginal discs has been shown to be clathrin and dynamin-dependent (Hemalatha et al, 2016). We suspect that the presence of these mutants may hamper the trafficking and nuclear localization of the C-terminal of DFz2, resulting in the phenotypes that we have described in this study. If this hypothesis holds true, it could be a good example of how mutant forms of clathrin disrupt signalling pathways (such as the Wnt signalling pathway) that are essential for proper neuronal development and function.

Overall, this study demonstrates that mutant forms of clathrin (L1047P and W1108R) disrupt clathrin-associated functions in *Drosophila,* leading to distinct behavioural phenotypes. Further, our work also predicts that patients carrying these mutations may show reduced survival. Our findings also establish *Drosophila* as a highly valuable genetic model for dissecting the cellular and molecular mechanisms underlying neurodevelopmental disorders. We propose that this system could be used to screen and identify potential therapeutic targets in a manner relevant to human physiology.

## Materials and methods

### Drosophila strains

Fly lines overexpressing either the wild-type (GFP-Chc) or the mutant forms (GFP-L1048P-Chc or GFP-W1109R-Chc) of *Drosophila* Chc under a UAS promoter and a GFP tag at the N-terminal were generated by WellGenetics (New Taipei City 221416, Taiwan). The overexpression constructs were injected into embryos and inserted into the attP2 site on chromosome 3L (68A4). Two individual lines for each genotype, described in Supplementary Table 1, were used in all the experiments. Other GAL4 driver lines used in the study have also been listed in Supplementary Table 1.

### Hemocyte isolation

Hemocytes were isolated by dissecting the third instar larvae under the dissecting microscope. Hemocytes were collected in S2 medium (Schneider’s medium) supplemented with 10% FBS. Hemocytes were then placed in a glass bottom 35mm dish and subjected to treatment or imaging.

### mBSA uptake assay

To prepare maleylated bovine serum albumin (mBSA), BSA (5mg) was dissolved in 1ml of carbonate buffer. 12.5mg of Maleic anhydride was added to 1 ml of BSA. pH was adjusted to 8.5 – 9. Maleylated BSA was dialyzed for 48 hours against 0.01M Ammonium bicarbonate and then for 16 hours against the carbonate buffer. The maleylated BSA was then tagged with an Alexa Fluor – 594 tag using Alexa Fluor™ 594 Microscale Protein Labeling Kit (Invitrogen, A30008) following manufacturer’s protocol.

For mBSA uptake, larvae were dissected and hemocytes were collected in serum-free Schneider’s Drosophila medium. Cells were incubated for 10–15 minutes at room temperature to allow them to settle down in glass-bottomed confocal dishes. Cells were incubated with mBSA at a concentration of 1 µg/ml for 5 mins and washed with ice-cold PBS. Fixation of hemocytes was done using 4% PFA for 20 minutes and cells were washed with 1x PBS before imaging.

### Site-directed mutagenesis

Human CHC17 cloned into a pEGFP-C1 vector backbone (GFP-CHC17KDP – gift from Stephen Royle; Addgene plasmid # 59799) was used as a template for site-directed mutagenesis. The mutations L1047P and W1108R were generated using the QuikChange II XL Site-Directed Mutagenesis Kit (Agilent Technologies, Catalog No. 200521) following the manufacturer’s instructions. The primer sequences used for SDM were:

L1047P-CHC-FP – 5’ gttatggagtatattaaccgc**cca**gataattat 3’

L1048PCHC-RP – 5’ atctggggcatcataattatc**tgg**gcggttaat 3’

W1108R-CHC-FP – 5’ cgttgcaatgaacctgcggtc**cga**agtcaactt 3’

W1108R-CHC-RP-5’ ctgggcttttgcaagttgact**tcg**gaccgcagg 3’

The positive clones carrying the point mutations (L1047P-CHC and W1108R-CHC) were confirmed by sequencing.

### Mammalian cell culture and transfection

HEK-293 cells were cultured on coverslips in DMEM (Gibco, catalog No. 11960-044) containing 10% FBS (Gibco, Catalog No. 10270-106), L-Glutamine (Gibco, catalog No. 25030-081), MEM Non-essential amino acids (Gibco, Catalog No. 11140-050), and 1X Penicillin/streptomycin (Gibco, catalog No.15070-063). Cells were grown in an incubator at 37°C and 5% (v/v) CO2.

For transfection, cells were seeded on 24 mm coverslips. 1.5µg of the desired plasmid was transfected using Lipofectamine 2000 reagent (Invitrogen, Catalog No. 11668019) as per the manufacturer’s instructions. The transfection mixture containing Lipofectamine and the plasmid was added dropwise to the cells. After 6 hrs, the media was replaced with fresh medium and the cells were kept in the incubator at 37°C. Imaging was done 48 hours after transfection. The wild-type CHC or either of the mutant forms of CHC (L1047P or W1108R) plasmids were used for transfection.

### Electrophysiology

All intracellular recordings were performed on wandering third instar larvae as described previously (Choudhury et al., 2016). Briefly, HL3 buffer containing 1.5 mM Ca^2+^ was used for larval dissection. Recordings were performed from muscle 6 of the A2 hemi-segment using sharp glass electrodes with a resistance of 20-30MΩ. mEJPs (miniature excitatory junction potentials) were recorded for 60 seconds, followed by recordings of EJPs (excitatory junction potentials) at 1Hz stimulation for 30 seconds. The high-frequency EJPs were recorded for 5 min by stimulating the nerves at 10 Hz. Excitatory junctional potentials (EJPs) were evoked by delivering stimulation pulses using a Grass S88 stimulator (Grass Instruments, Astro-Med,Inc.). The resulting signals were amplified with an Axoclamp 900A amplifier, digitized via a Digidata 1440A digitizer, and recorded using pClamp10 software (Molecular Devices). Only muscles with a resting membrane potential between −60 and −75 mV were included in the analysis. The data were analyzed using the Mini Analysis program (Synaptosoft, Decatur,GA).

### Antibodies and Immunostaining

For staining of larval NMJs, wandering third instar larvae were dissected and fixed with 4% PFA. After fixation, they were permeabilized with 0.3% Triton-X-100 and blocked with 5% NGS, followed by overnight incubation with the primary antibody at 4°C. This was followed by washing and incubation with the secondary antibody along with Phalloidin and HRP for 4 hours at room temperature. The larvae were then mounted onto glass slides and imaged. For staining of adult fly brains, a similar protocol was followed. The antibodies used in the study have been described in Supplementary Table 2.

### Proboscis extension response (PER) assay

The protocol was adapted from (Masek et al., 2015). 5-10 days old males and females were collected and starved of food for 24 hours by placing them in a vial with wet filter paper. The flies were then anesthetised on ice and stuck onto glass slides on their thorax and wing base using nail polish. They were then allowed to recover for 3 hours prior to the experiment.

Flies were satiated with water before and during the experiment. Flies that did not initially satiate within 5 minutes were excluded from conditioning. In the pre-test, each fly was given 100 mM sucrose on their tarsi three times with 10 sec inter-trial interval and the number of full proboscis extensions was recorded. Any fly that did not take out its proboscis all three times was also excluded from the training. During training, a droplet of 50mM denatonium benzoate (Sigma, catalog No. 3734-33-6) was placed on the extended proboscis and flies were allowed to drink it for up to 2 seconds or until they retracted their proboscis. After each session, tarsi and proboscis were washed with water and flies were allowed to drink to satiation. There were three trials with a 10-minute interval between each trial, followed by testing, 10 minutes after the third trial. During each trial, the fly was presented with sucrose thrice with a 10-second inter-interval.

### Locomotion assay

To study larval locomotion, wandering third instar larvae were collected and placed on a plate with 1% agar. They were allowed to move along the plate and 120 second videos were recorded. The locomotion behaviour was then quantified using the Tracker video analysis and modeling Tool, open source physics, available online.

To study locomotion in adults, the negative geotaxis assay was performed. 5-10 days old males and females were placed in a 50ml measuring cylinder. After 15 minutes of acclimatization, they were tapped down to the bottom of the cylinder and allowed to climb. 90-second videos were recorded and analysed manually. Time taken for 50% of the flies to cross the 10cm mark for each genotype was plotted.

### Survival and eclosion percentage calculations

All flies eclosing on the same day were collected in food vials and the vials were flipped every 5 days. The males and females were kept separately. Kaplan-Meier curves were plotted to study differences in survival between genotypes.

In order to look at eclosion numbers, 15 larvae of each genotype were placed in a food vial and the percentage of adults eclosing from the pupae were plotted.

### Confocal imaging and image analysis

For imaging of larval NMJs, hemisegment A2/A3 at the 6^th^ and 7^th^ muscle of the third instar larva was imaged. All images of the larval NMJs and adult fly brains were acquired using Olympus FV3000 confocal laser scanning microscope at either 40x or 60x.

Timelapse imaging of hemocytes and HEK was performed along a single X-Y plane close to the glass cover slip to look at clathrin dynamics. Imaging was carried out for 5 minutes at 5 second intervals using a Nikon Ti Eclipse confocal microscope at 100x.

The mean instantaneous velocities of clathrin-coated structures (CCSs) were quantified by calculating the distance moved by an individual CCS for each 5 sec interval of the timelapse images, and averaging these instantaneous velocities for that CCS.

For mBSA uptake experiments, mBSA intensity in each cell was calculated using ImageJ. For the calculation of spectrin intensity, ROI was selected manually by marking the boutons in the HRP channel and calculating the intensity in the spectrin channel following background subtraction. For experiments where fluorescence intensity was quantified, imaging for each experiment was done using the same laser power.

Branch number, total number of boutons/NMJ, number of NC82 puncta and the number of Dlg-negative boutons (HRP +ve and Dlg -ve) were counted manually. Calculation of the average bouton area was done using ImageJ.

## Acknowledgements

This work was supported by funds to DS and AM from Department of Biotechnology (BT/PR43190/MED/97/573/2021); CRG/2021/00599 from SERB to VK; Senior Research Fellowship from UGC to JD. We would like to thank Dr. Gaurav Das, and Radhika Mohandasan at NCCS Pune for their help in conducting the learning and memory experiments in the study. We thank members of the Subramanyam lab for inputs and constructive discussion. DS thanks AVR for support.

## Author contributions

**Jyoti Das –** conceptualization; data curation and analysis – eclosion percentage (Fig. 1A,B; Supplemental Fig.1C), Site directed mutagenesis for vesicle dynamics experiment (Supplemental Fig. 1F, G) ligand uptake experiment (Fig.1F-I; supplemental Fig. 1J,K), neuronal morphology at the NMJ (Fig.3A-D; Supplemental Fig.3 A-D), learning and memory (Fig.5B-D; Supplemental Fig. 5A,B), active zone and post-synaptic markers at the NMJ (Fig.3E, F; Fig.4A-H; Supplemental Fig. 3E,F; Supplemental Fig. 4A-H); data visualization, Writing – original draft, review and editing.

**Manisha Datta** – data curation and analysis – survival (Fig. 1C, D; Supplemental Fig. D, E), larval locomotion (Fig.2E, F; Supplemental Fig. 2E), Writing – review and editing.

**Srikanth Pippadpally** – data curation and analysis – electrophysiology experiments (Fig. 3A-D; Supplemental Fig. 3A-D); Writing – review and editing.

**Bhavika Jethani** – data curation and analysis – ligand uptake (Fig. 1F-I; supplemental Fig. 1J,K), active zone and post-synaptic markers at the NMJ (Fig.3E, F; Fig.4A-H; Supplemental Fig. 3E, F; Supplemental Fig. 4A-H), Writing – review and editing.

**Surya Bansi Singh** – data curation, analysis and visualization – vesicle dynamics experiments (Fig. 1E; Supplemental Fig. 1H, I); Writing – review and editing.

**Akankhya Jena** – data curation for learning and memory experiment (Fig.5B, C; Supplemental Fig. 5A, B), data analysis for neuronal morphology (Fig.3A-D; Supplemental Fig.3 A-D).

**Vimlesh Kumar** – data curation and analysis – electrophysiology experiments (Fig. 3A-D; Supplemental Fig. 3A-D), Writing – review and editing, funding acquisition

**Amitabha Majumdar** – Conceptualization, Writing – review and editing, funding acquisition.

**Deepa Subramanyam** – Conceptualization, data visualization, Writing – original draft, review and editing, funding acquisition, supervision and project administration.

**Disclosure:** The authors declare no conflict of interest.

---

**Supplementary Figure1: A)** Sequence alignment of *Homo sapiens* and *Drosophila melanogaster* clathrin heavy chain protein. The boxed residues represent L1047P,which represents leucine mutated to proline and W1108R, which represents tryptophan mutated to arginine. **B)** Schematic representation of the construct used to generate flies overexpressing wild type or mutant forms of Chc. The Chc sequence along with GFP at the N terminal was cloned into the pUAST vector and introduced in *Drosophila* at the attp2 site on chromosome 3L (68A4). **C)** Eclosion percentage of *Elav>GFP-L1048P-Chc* Line2 and *Elav>GFP-W1109R-Chc* Line2 compared to *Elav>GFP-Chc* Line2. n=60; N=4. P-value calculated using Kruskal-Wallis test for non-parametric data. *p<0.05. **D)** Kaplan-Meier curve depicting survival of *Tub>GFP-W1109R-Chc* Line2 (N=36) flies compared to *Tub>GFP-Chc* Line2 (N=43), *UAS-GFP-W1109R-Chc/+* Line2 (N=49), *UAS-GFP-Chc/+* Line2 (N=49) and *CsBz* (N=44) controls. **E)** Kaplan-Meier curve depicting survival of *Elav>GFP-W1109R-Chc* Line2 (N=37) and *Elav>GFP-L1048P-Chc* Line2 (N=32) flies compared to Elav>GFP-Chc Line2 (N=40), *UAS-GFP-W1109R-Chc/+* Line2 (N=49), *UAS-GFP-L1048P-Chc/+* Line2 (N=50) *UAS-GFP-Chc/+* (N=49) and *CsBz* (N=44) controls. **F)** Chromatogram showing site directed mutagenesis performed to generate the L1047P CHC mutant in human CHC cloned in pEGFP-C1 vector. **G)** Chromatogram showing site directed mutagenesis done to generate the W1108R CHC mutant in human CHC cloned in pEGFP-C1 vector. **H)** Representative images of *GFP-Chc17*, *GFP-L1047P-Chc* and *GFP-W1108R-Chc* in HEK293 cells. Scale bar-5µm. **I)** Graph representing line speed of vesicles in HEK293 cells expressing *GFP-Chc* (n=279), and *GFP-W1108R-Chc* (n=582). P-value calculated using Kruskal-Wallis test for non-parametric data. ****P≤0.0001. **J)** Representative images of mBSA uptake in *GFP-Chc* Line2, *GFP-L1048P-Chc* Line2 and GFP-W1109R-Chc Line2 expressing hemocytes. Ligand uptake was performed at 37°C. Scale bar-5µm. **K)** Graph quantifying the mBSA intensity inside hemocytes expressing *GFP-Chc* Line2 (n=44), *GFP-L1048P-Chc* Line2 (n=35) and *GFP-W1109R-Chc* Line2 (n=41). Ligand uptake was performed at 37°C. Statistical significance calculated using the Kruskal-Wallis test for non-parametric data. All data is represented as mean ± SEM.

**Supplementary Figure 2: A)** Histograms showing average mEJP frequency in *Elav-Gal4>UAS-GFP* (2.93 ± 0.30), *Elav>GFP-Chc* Line2 (2.21 ± 0.12), *Elav>GFP-L1048P-Chc* Line2 (2.32 ± 0.14) and *Elav>GFP-W1109R-Chc* Line2 (2.36 ± 0.15). Statistical analysis is based on one-way ANOVA followed by post-hoc Tukey’s multiple-comparison test. **B)** Histograms showing average mEJP amplitude in *Elav-Gal4>UAS-GFP* (0.87 ± 0.04), *Elav>GFP-Chc* Line2 (1.03 ± 0.07), *Elav>GFP-L1048P-Chc* Line2 (0.86 ± 0.07) and *Elav>GFP-W1109R-Chc* Line2 (0.99 ± 0.05). Statistical analysis is based on one-way ANOVA followed by post-hoc Tukey’s multiple-comparison test. **C)** Histograms showing average EJP amplitude in *Elav-Gal4>UAS-GFP* (46.8 ± 1.88), *Elav>GFP-Chc* Line2 (44.1 ± 1.14), *Elav>GFP-L1048P-Chc* Line2 (42.7 ± 1.37) and *Elav>GFP-W1109R-Chc* Line2 (42.4 ± 0.49). Statistical analysis is based on one-way ANOVA followed by post-hoc Tukey’s multiple-comparison test. **D)** Histograms showing quantal content in *Elav>GFP* (50.2 ± 4.45), *Elav>GFP-Chc* Line2 (44.7 ± 3.69), *Elav>GFP-L1048P-Chc* Line2 (51.8 ± 5.18), and *Elav>GFP-W1109R-Chc* Line2 (43.5 ± 2.32). **E)**Graph representing the distance travelled by *Elav>GFP-W1109R-Chc* Line2, *Elav>GFP-L1048P-Chc* Line2 larvae compared to controls *Elav>GFP-Chc* Line2, *UAS-GFP-W1109R-Chc/+* Line2, *UAS-GFP-L1048P-Chc/+* Line2, *UAS-GFP-Chc/+* Line2 and *Elav-GAL4/+*. Statistical significance calculated using one-way ANOVA. *P≤0.05; **P≤0.005; ***P≤0.0005. 10 animals of each genotype were studied. The larvae were kept at 37°C for 20 minutes. All data is represented as mean ± SEM.

**Supplementary Figure 3: A)** Representative images of neuronal morphology in *Elav>GFP-Chc* Line2, *Elav>GFP-L1048P-Chc* Line2 and *Elav>GFP-W1109R-Chc* Line2 larvae. Neurons at the A2/A3 hemisegment on the 6^th^/7^th^ muscle were imaged. 5-7 animals were imaged for each genotype. Scale bar – 20µm. **B)** Graph representing total bouton number in *Elav>GFP-Chc* Line2, *Elav>GFP-L1048P-Chc* Line2 and *Elav>GFP-W1109R-Chc* Line2 larvae. Statistical significance calculated using the Kruskal-Wallis test. **C)** Graph representing average bouton size in *Elav>GFP-Chc* Line2, *Elav>GFP-L1048P-Chc* Line2 and *Elav>GFP-W1109R-Chc* Line2 larvae. Statistical significance calculated using the Kruskal-Wallis test. **D)** Graph representing total branch number in *Elav>GFP-Chc* Line2, *Elav>GFP-L1048P-Chc* Line2 and *Elav>GFP-W1109R-Chc* Line2 larvae. Statistical significance calculated using the Kruskal-Wallis test. **E)** Representative images showing the active zones in *Elav>GFP-Chc* Line2, *Elav>GFP-L1048P-Chc* Line2 and *Elav>GFP-W1109R-Chc* Line2 larvae. Scale Bar – 10µm. **F)** Graph representing the number of active zones per bouton in *Elav>GFP-Chc* Line2, *Elav>GFP-L1048P-Chc* Line2 and *Elav>GFP-W1109R-Chc* Line2 larvae. 40-90 boutons were counted for each genotype. Statistical significance calculated using the Kruskal-Wallis test. All data is represented as mean ± SEM.

**Supplementary Figure 4: A)** Representative images showing the Dlg positive and negative boutons in *Mef2>GFP-Chc* Line2, *Mef2>GFP-L1048P-Chc* Line2 and *Mef2>GFP-W1109R-Chc* Line2 larvae. Scale bar-10µm. **B)** Graph quantifying the number of Dlg negative boutons in *Mef2>GFP-Chc* Line2, *Mef2>GFP-L1048P-Chc* Line2 and *Mef2>GFP-W1109R-Chc* Line2 larvae. 8-10 animals were dissected for each genotype. Statistical significance calculated using the Kruskal-Wallis test. **P≤0.005; *** P≤0.0005. **C)** Representative images showing Spectrin in *Mef2>GFP-Chc* Line2, *Mef2>GFP-L1048P-Chc* Line2 and *Mef2>GFP-W1109R-Chc* Line2 larvae. Scale bar-10µm. **D)** Graph quantifying spectrin intensity in *Mef2>GFP-Chc* Line2, *Mef2>GFP-L1048P-Chc* Line2 and *Mef2>GFP-W1109R-Chc* Line2 larvae. 8-10 animals were dissected for each genotype. Statistical significance calculated using the Kruskal-Wallis test. **** P≤0.0001. **E)** Representative images showing the Dlg positive and negative boutons in *Elav>GFP-Chc* Line2, *Elav>GFP-L1048P-Chc* Line2 and *Elav>GFP-W1109R-Chc* Line2 larvae. Scale bar-5µm. **F)** Graph quantifying the number of Dlg negative boutons in *Elav>GFP-Chc* Line2, *Elav>GFP-L1048P-Chc* Line2 and Elav>GFP-W1109R-Chc Line2 larvae. 8-10 animals were dissected for each genotype. Statistical significance calculated using the Kruskal-Wallis test. *P≤0.05; *** P≤0.0005. **G)** Representative images showing Spectrin in *Elav>GFP-Chc* Line2, *Elav>GFP-L1048P-Chc* Line2 and *Elav>GFP-W1109R-Chc* Line2 larvae. Scale bar- 10µm. **H)** Graph quantifying spectrin intensity in *Elav>GFP-Chc* Line2, *Elav>GFP-L1048P-Chc* Line2 and *Elav>GFP-W1109R-Chc* Line2 larvae. 8-10 animals were dissected for each genotype. Statistical significance calculated using the Kruskal-Wallis test. **** P≤0.0001. All Data is represented as Mean ± SEM.

**Supplementary Figure 5: A)** PER assay for *Tub>GFP-W1109R-Chc* Line2 flies and *Tub>GFP-Chc* Line2 and parental controls. N=10-20 for each genotype. All flies were 3-10 days old post eclosion. Statistical significance was calculated using a two-way ANOVA followed by Dunnett multiple comparison method. *P≤0.05. Statistical significance shown using *Tub>GFP-Chc* Line2 as a control. **B)** PER assay for *Elav>GFP-L1048P-Chc* Line2, *Elav>GFP-W1109R-Chc* Line2 flies and *Elav>GFP-Chc* Line2 and parental controls. N=10-20 for each genotype. All flies were 3-10 days old post eclosion. Statistical significance was calculated using a two-way ANOVA followed by Dunnett multiple comparison method. *P≤0.05; **P≤0.005. Statistical significance shown using *Elav>GFP-Chc* Line2 as a control.

## Supplementary Tables

**Supplementary Table 1:** Fly lines used in this study.

| Fly Line | Source | Stock Number (If available) |
| --- | --- | --- |
| *CsBz* | Stock, Dr. Amitabha Majumdar, NCCS | -- |
| *Tub-GAL4* | Stock, Dr. Gaurav Das, NCCS | -- |
| *Mef2-GAL4* | BDSC | 27390 |
| *Cg-GAL4* | BDSC | 7011 |
| *Elav-GAL4* | BDSC | 458 |
| *UAS-GFP-Chc Line1* | WellGenetics (this study) | SWGA - 7205 |
| *UAS-GFP-Chc Line2* | WellGenetics (this study) | SWGA- 7206 |
| *UAS-GFP-L1048P-Chc Line1* | WellGenetics (this study) | SWGA - 6957 |
| *UAS-GFP-L1048P-Chc Line2* | WellGenetics (this study) | SWGA- 6956 |
| *UAS-GFP-W1109R-Chc Line1* | WellGenetics (this study) | SWGA- 6959 |
| *UAS-GFP-W1109R-Chc Line2* | WellGenetics (this study) | SWGA- 6958 |

**Supplementary Table 2:** Antibodies used in this study.

| Antibody | Company (Catalogue No.) | Dilution (I.F.) |
| --- | --- | --- |
| Phalloidin-488 | Invitrogen (A12379) | 1:250 |
| HRP-555 | Jackson<br>ImmunoResearch<br>(123-295-021) | 1:250 |
| HRP-647 | Jackson<br>ImmunoResearch<br>(123-605-021) | 1:250 |
| nc82 | DSHB (AB_2314866) | 1:10 |
| Discs Large (Dlg) | DSHB (AB_528203) | 1:50 |
| Spectrin | DSHB (AB_528473) | 1:50 |
| FasII | DSHB (AB_528235) | 1:50 |
| Alexa Flour 647<br>goat anti-mouse | A-21235 | 1:500 |

