## Supplementary figures and images for "Pathogenic mutations in Clathrin Heavy Chain associated with intellectual disability impair synaptic architecture and learning in *Drosophila*"

### Supplemental Figure 1

#### Supplemental figure 1

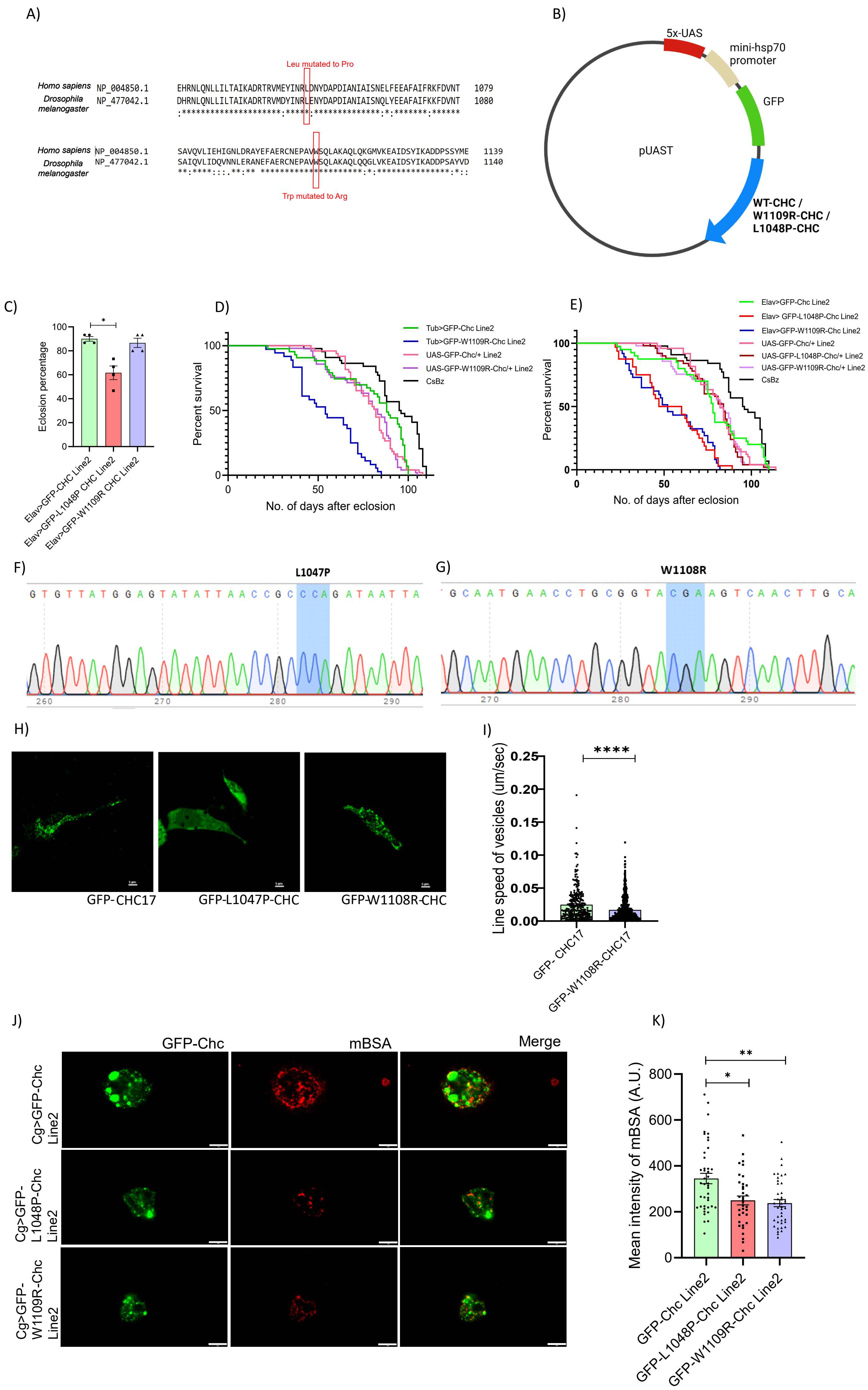

### Supplemental Figure 2

#### Supplemental figure 2

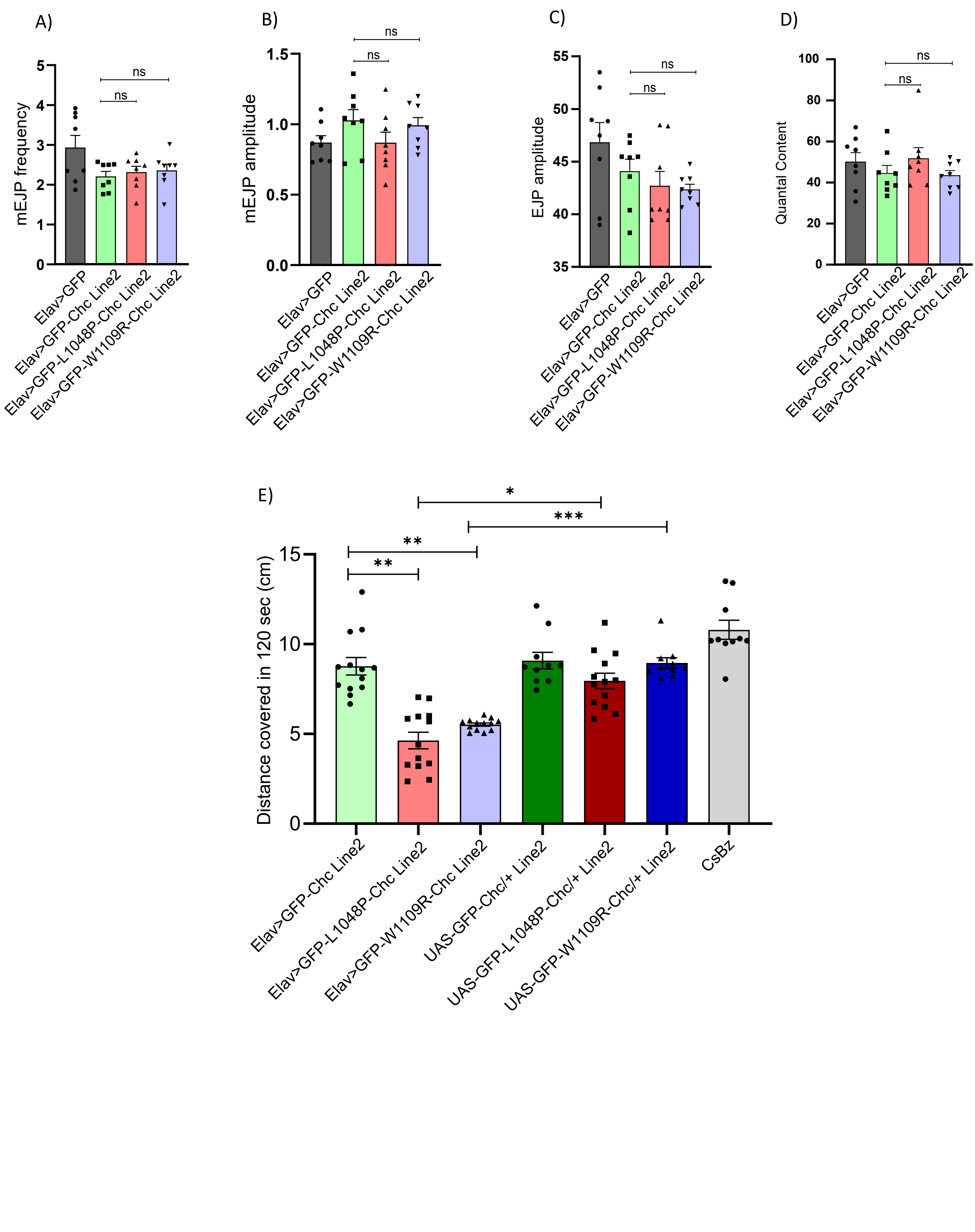

### Supplemental Figure 3

#### Supplemental figure 3

A)

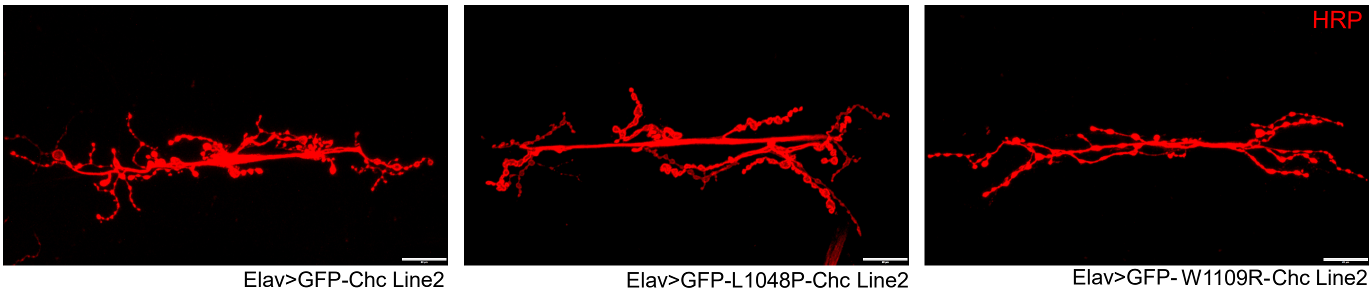


B)

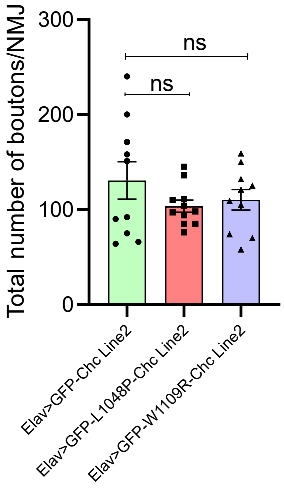


C)

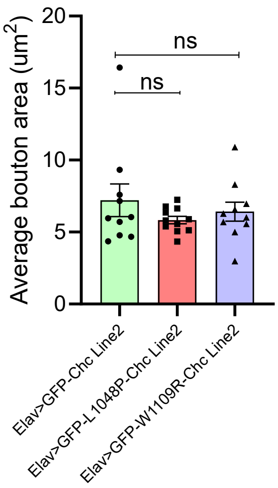


D)

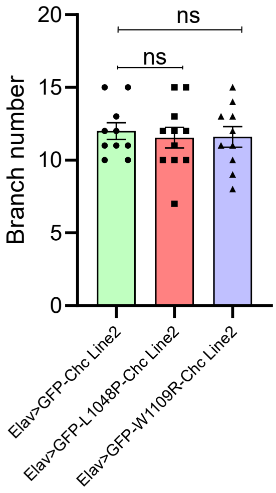


E)

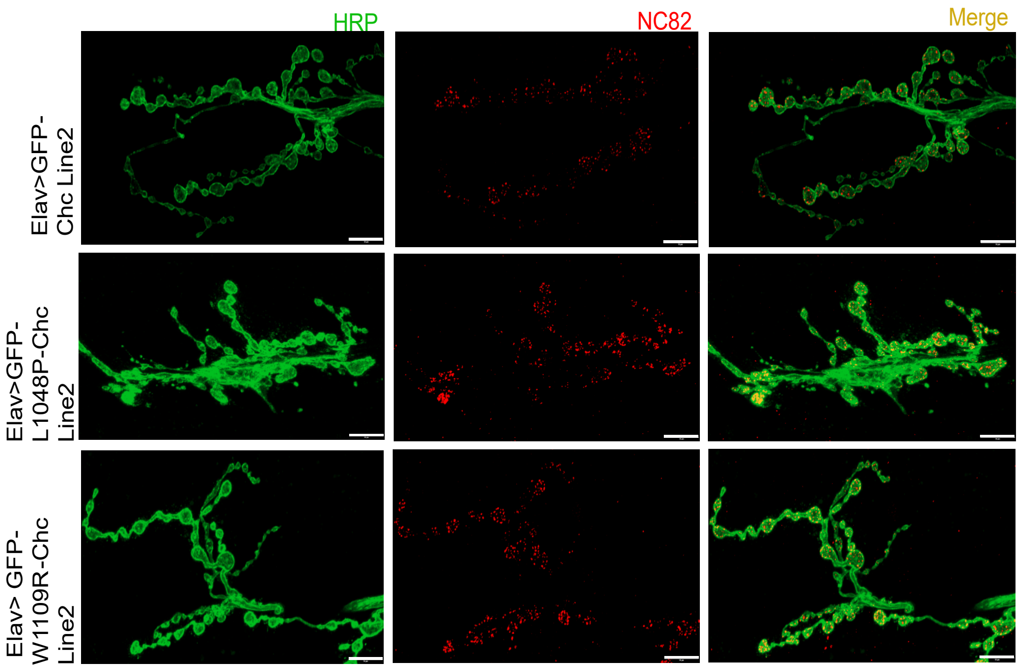


F)

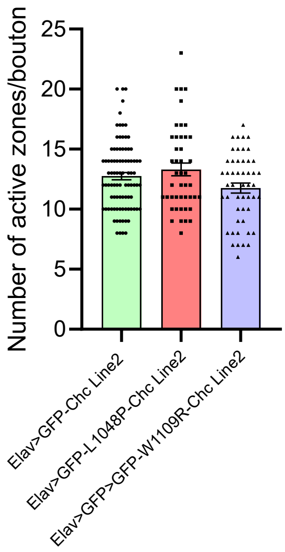

### Supplemental Figure 4

#### Supplemental figure 4

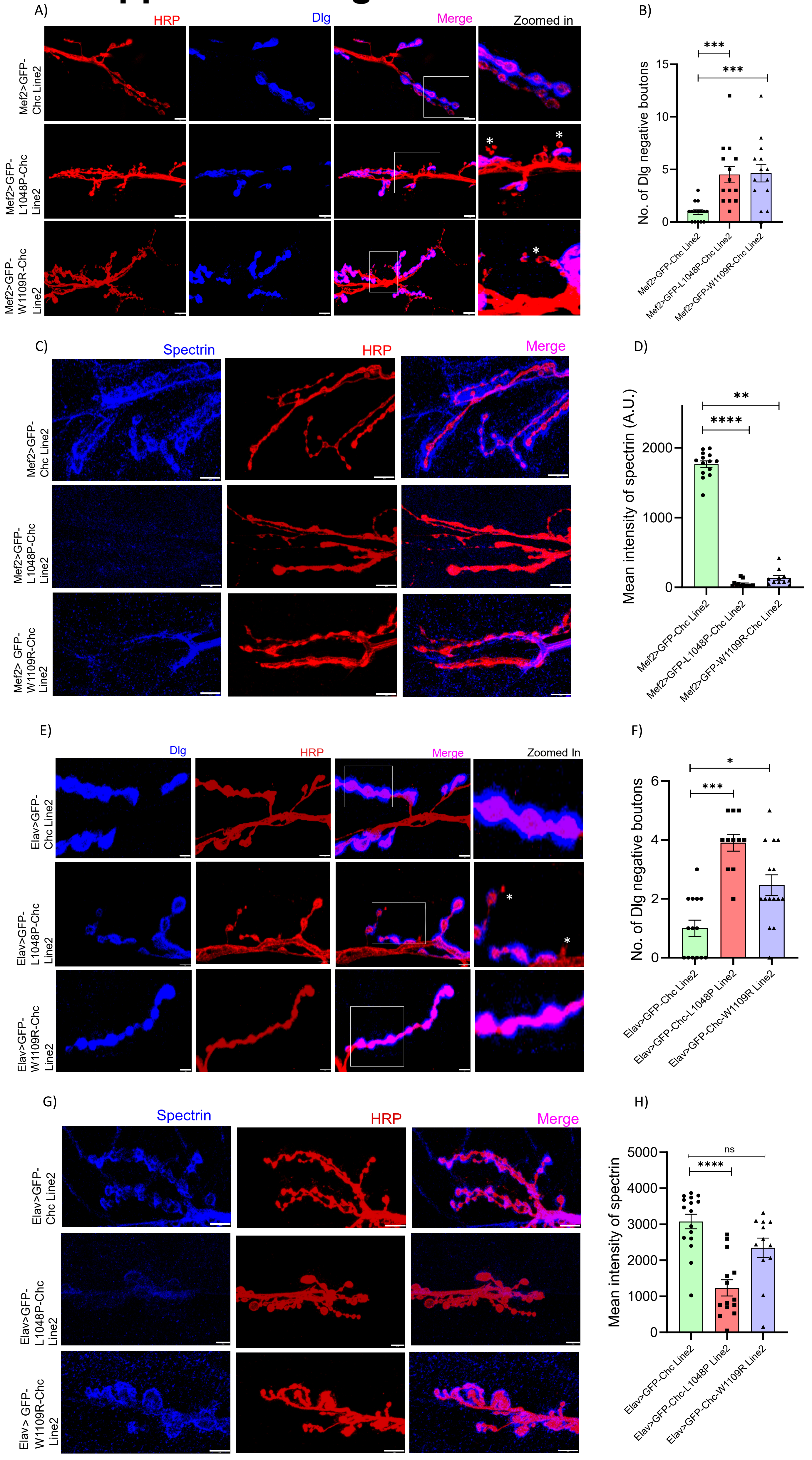

### Supplemental Figure 5

#### Supplemental figure 5

A)

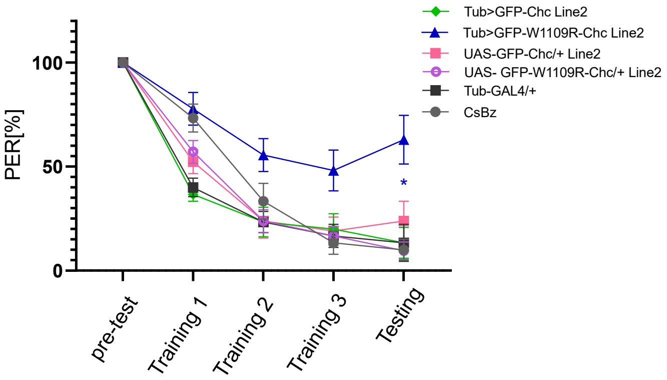


B)

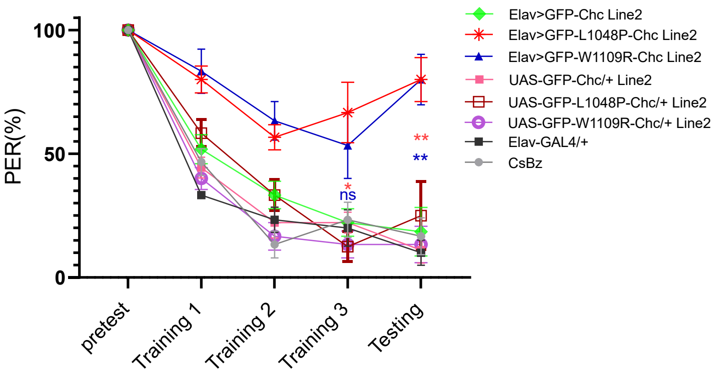
